# Live poultry feeding and trading network and the transmission of avian influenza A(H5N6) virus in a large city in China, 2014-2015

**DOI:** 10.1101/2020.10.21.349787

**Authors:** Rusheng Zhang, Zhao Lei, Chan Liu, Yuanzhao Zhu, Jingfang Chen, Dong Yao, Xinhua Ou, Wen Ye, Zheng Huang, Li Luo, Biancheng Sun, Tianmu Chen

**Affiliations:** Changsha Center for Disease Control and Prevention, Changsha, Hunan Province, People’s Republic of China; State Key Laboratory of Molecular Vaccinology and Molecular Diagnostics, School of Public Health, Xiamen University, Xiamen, Fujian Province, People’s Republic of China

**Author notes:** Correspondence: Tianmu Chen, State Key Laboratory of Molecular Vaccinology and Molecular Diagnostics, School of Public Health, Xiamen University, 4221-117 South Xiang’an Road, Xiang’an District, Xiamen, Fujian Province, People’s Republic of China. These authors contributed equally to this study (Rusheng Zhang, Zhao Lei, Chan Liu).

**Keywords:** H5N6, avian influenza virus, transmission, live poultry market, network

## Abstract

To understand clearly the mechanism of H5N6 transmission, the role of a live poultry feeding and trading network should be explored deeply. However, there is little data to show the network in an area. In this study, we performed a field epidemiological investigation to collect the numbers of farms, wholesale markets, and live poultry retail markets, and the numbers of purchased (from where) and sold (to where) live birds in Changsha City, China in 2014. We also collected samples from the network in the city from January 2014 to March 2015, including the LPMs visited by the patient known to be infected with A(H5N6) virus, and sequenced the genomes of 10 A(H5N6) viral strains isolated from these environmental samples, to determine the source of virus that infected the patient reported in Changsha. Additionally, we collected and analyzed A(H5N6) virus genome sequence isolated from humans, poultry, and LPMs registered in National Center for Biotechnology Information (NCBI; NIH, Bethesda, MD, USA) 14, to determine the source of other human infections with A(H5N6) virus reported in China. Changsha City in China has a large live poultry feeding and trading network (LPFTN) which had 665 farms, 5 wholesale markets, and 223 live poultry retail markets in 2014. The network covered nine provinces and purchased and sold more than 150000 live birds every day. About 840 environmental samples collected from the LPFTN network from January 2014 to March 2015. About 8.45% (71/840) environmental samples were shown to be positive for N6 and 10 full genome sequences of H5N6 virus were analyzed. We performed phylogenetic analyses and virus characterization, which demonstrated that the H5N6 viral strains isolated from Chinese patients were closely related to those isolated from the poultry and environmental samples obtained from the LPFTN network. This indicates that the network with a large volume of live poultry provides a platform for the transmission of H5N6, and provides an infectious pool which makes the people in high risk.

**Importance:** Avian influenza A(H5N6) virus is an emerging threat to public health. With several human cases reported in recent years, it has become a dominant avian influenza virus (AIV) subtype in China. Live poultry feeding and trading network (LPFTN) contributes to the presence of novel AIV. By a field epidemiological investigation in the complex LPFTN in Changsha, we demonstrated that the H5N6 viral strains isolated from Chinese patients were closely related to those isolated from the poultry and environmental samples obtained from the network, suggesting the LPFTN with a large volume of live poultry provides a platform for the transmission of H5N6 and creates an infectious pool which makes people in high risk. Considering the wide circulation and dynamic reassortment of this highly pathogenic avian influenza H5N6 virus, it should be carefully monitored in poultry and humans due to the pandemic potential.

## Introduction

As of December 1, 2016, a total of 17 laboratory-confirmed cases of human infection with avian influenza A(H5N6) virus has been detected in Sichuan, Guangdong, Yunnan, Hunan, Hubei, Guangxi, and Anhui provinces of China since 2014, which resulted in 6 deaths (Table 1) ^1^. With the exception of 4 cases where the source of exposure was unknown, in all other cases, the individuals were shown to be previously exposed to poultry or had access to the live poultry markets (LPMs), but the specific sources of infection have not been clearly identified^2-5^.

**Table 1.**
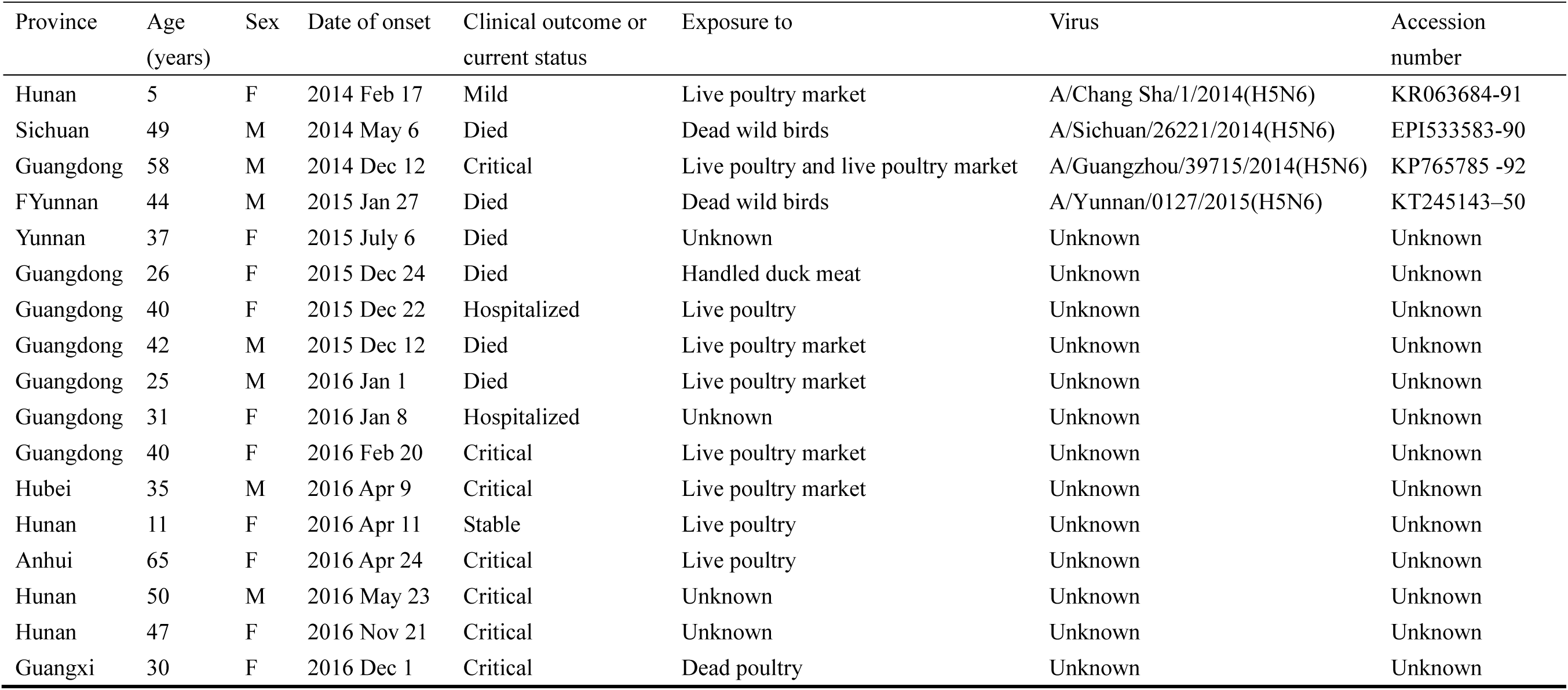
Patients infected with avian influenza A(H5N6) virus, from February 17, 2014 through December 1, 2016

In the early 2015, Changsha Center for Disease Control and Prevention (CDC) reported that one patient had been infected with A(H5N6) virus, which was confirmed by genome sequencing. Epidemiological investigation determined that one LPM might have represented the source of exposure, but further analyses did not identify the virus in the LPMs, and the source of this infection could not be determined^6^.

In December 2013, one A(H5N6) viral strain was isolated from a sewage sample in a LPM in Jiangsu, China ^7^, while in March 2014, in Laos, poultry infection with the same virus was reported, and shown to have originated due to the reassortment of A(H5N1) and A(H6N6) viruses found in duck populations in southern and eastern China ^8^. During the same month, ducks in Guangdong province in southern China were shown to be infected with A(H5N6) virus^9^. Genetic and phylogenetic analyses detected these viruses at the LPMs in China in 2014, which had evolved into two distinct lineages, Sichuan and Jiangxi^10^. Furthermore, LPMs have previously been suggested to represent a major risk factor for the infection of humans with the avian A influenza viruses, such as A(H5N1), A(H7N9), and A(H10N8) viruses^11-13^, but their role in the development of human A(H5N6) viral infections has not been firmly established.

The LPMs are parts of live poultry feeding and trading network (LPFTN). To understand clearly the mechanism of H5N6 transmission, the role of the LPFTN network should be explored deeply. However, there is little data to show the network in an area.

In this study, we performed a field epidemiological investigation to collect the numbers of farms, wholesale markets, and live poultry retail markets, and the numbers of purchased (from where) and sold (to where) live birds in Changsha City, China in 2014. We also collected samples from the LPFTN network in the city from January 2014 to March 2015, including the LPMs visited by the patient known to be infected with A(H5N6) virus, and sequenced the genomes of 10 A(H5N6) viral strains isolated from these environmental samples, to determine the source of virus that infected the patient reported in Changsha. Additionally, we collected and analyzed A(H5N6) virus genome sequence isolated from humans, poultry, and LPMs registered in National Center for Biotechnology Information (NCBI; NIH, Bethesda, MD, USA) ^14^, to determine the source of other human infections with A(H5N6) virus reported in China.

## Materials and methods

### Study area

Changsha (27°51’∼28°41’ N, 111°53’∼114°15’ E), a large city with 7.04 million people in central south China, is the capital of Hunan Province. It includes 9 sub-areas (6 districts, 2 counties, and 1 county-level city), with a total of 4,586 hospitals, clinics, and public health departments all over the city. Live poultry consumption is the main consumption mode of poultry meat in the city.

### Investigation on live poultry feeding and trading network

To understand the live poultry feeding and trading network (LPFTN) in the city, we collected data on all farming sites, wholesale markets and retail markets in Changsha from June to July 2014. The field epidemiological investigation was performed by Changsha Animal Disease Control Center. The information of breeding sites, which collected from poultry farms and individual farmers who raise more than 100 live birds, includes: basic information of farmers (address, area of the breeding site, number of farmers, and information of routine cleaning and disinfection), information of live poultry breeding and immunization (number of stock poultry, number of annual sales, the place where the young poultry purchased, and information of vaccination), and live poultry trading (sales place, and number of sold poultry). The information collected in live poultry wholesale market includes: basic market information (name, address, area of the market, and information of routine cleaning and disinfection), the number of wholesalers, basic information of each wholesalers (area and number of workers), live poultry purchasing situation (type of poultry, inventory, purchasing volume of poultry, daily average sales volume, and purchasing source), sales situation (place where the poultry sold to, and daily number of sold poultry). The information collected in live poultry retail market includes: basic market information (name of the market, address, area of the market, information of routine cleaning and disinfection, and total number of stalls), stall information (area of the stall, and number of workers), live poultry stock situation (type of poultry, daily number of the poultry remained in the stall, daily average sales volume, and source of the poultry).

### Monitoring data of environmental specimens

From January 2014 to March 2015, the environmental samples of live poultry farming and trading places in Changsha City were collected systematically, covering all districts and counties (cities) of the city. The types of monitoring sites include farms, wholesale markets and retail markets. The environmental specimens collected include: sewage polluted by poultry (including water for cleaning slaughtered poultry, processing poultry tools, and cages), poultry manure (including chicken, duck, pigeon manure), poultry cages (doors) surface smearing, poultry falling feathers. The sewage mainly consists of water samples contaminated by poultry in ditches or pools beside poultry activities or cages. 15-20 ml of each sample is collected and placed in a tube with an outer spiral cover of 50 ml. The poultry excrement was collected by cotton swab for 5 g and placed in the sampling tube. Avian feathers were collected with tweezers and placed in a sampling tube. The surface of the cage (door) is wiped with a swab with a polypropylene fiber head contaminated with the sample solution at three to five different parts of the cage surface. Then the wiped swab is put into a sampling tube containing 5 ml of Hank’s sample solution, and the tail is discarded. At the same time, registration and collection time, the district, county, market name and other information. The collected samples were sent to the influenza surveillance network laboratory of Changsha CDC for nucleic acid detection of avian influenza virus within 24 hours. All samples were stored in the VTM Transportation Medium (Qingdao Hope Bio-Technology Co., Ltd, Qingdao, Shandong, China), which contained Hank’s balanced salt solution with 0.5% bovine serum albumin and antibiotics, at −80°C until further processing.

### Surveillance of human cases

There are two surveillance systems (influenza surveillance system, ISS and pneumonia surveillance system, PSS) to monitor the virus in the city. The ISS system, based on two sentinel hospitals which are both tertiary hospitals, was set up in Changsha in September 2005. In 2008, the ISS system became a branch of national influenza surveillance network. The system monitors the virus based on influenza-like illness (ILI). Throat swab specimens of ILI cases were collected for testing the influenza virus by reverse transcription polymerase chain reaction (RT-PCR) and / or cell culture in the laboratory of Changsha CDC. The PSS was built in Changsha in March, 2009. Pneumonia related inpatient departments of in 49 hospitals in Changsha were enrolled into the system. This system monitors pneumonia cases among inpatient population. When cases were suspected as avian influenza infection, their throat swabs or low respiratory tract specimens would be collected to detect the influenza viruses by RT-PCR. In this study, we collected the data of the two systems from January to December, 2014.

### RNA extraction and the detection of influenza virus subtype N6

Viral RNAs were extracted from 1,240 environmental samples using the QIAmp Viral RNA Mini Kit (Qiagen, Germany), according to the manufacturer’s instructions. The extracted RNA samples were eluted in 60 μL of nuclease-free water and stored at −80°C until further analyses. Each sample was screened using the conventional reverse transcription PCR (RT-PCR) for influenza typing using primers specific to N6, as previously described^15^.

### Genome sequencing and phylogenetic analysis

A total of eight universal primer sets were used to amplify the full genome of influenza A virus, which is composed of eight segments ^15^, followed by sequencing via SuperScript III One-Step RT-PCR System with Platinum *Taq* RT-PCR (Life Technologies, USA) conducted by Life Technologies Biotechnology (Shanghai, China). Full genome sequences of 10 avian influenza A(H5N6) viruses detected in the environmental LPM samples were deposited in GenBank (accession numbers: MH156460-MH156539).

The 10 environmental A(H5N6) virus genome data obtained in current study were characterized and phylogenetically analyzed with the corresponding gene segments of A(H5N6) virus obtained from GenBank, including the poultry and environmental samples in East and Southeast Asia, as well as four available samples from H5N6-infected patients reported by WHO (GenBank accession numbers: KR063684-91, KP765785-92, KT245143-50, and EPI533583-90). Phylogenetic trees were constructed in Molecular Evolutionary Genetics Analysis (MEGA) software version 5.2 with the neighbor-joining method using the Tamura-Nei model to calculate the genetic distances between sequences, and bootstrap resampling was done for 1000 repeats to estimate the reliability of the phylogenetic tree. The nucleotide sequences were then translated into amino acid protein sequences to detect the mutations.

## Results

### The LPFTN network

According to the field epidemiological investigation, there are 665 farms in Changsha, among which the smallest stock of live birds is 100, the largest is 3000000, the average stock is 12538, and the cumulative stock is 8212668. The smallest sales volume was 90, the largest was 12000000, the average sales volume of farms was 76142, and the cumulative sales volume was 34644815.

There are 5 wholesale markets and 162 wholesalers in Changsha. Among them, the smallest stock of live birds is 0, the largest is 640, the average stock is 28, the cumulative stock is 4525. The daily purchasing and selling volume was 1311444.

There are 223 live poultry retail markets and 493 retailers in Changsha. Among them, the smallest stock of live birds is 0 (i.e. the daily stock is sold out), the largest is 1800, the average stock is 62, and the cumulative stock is 30318. The daily purchasing and selling volume is 152051.

Overall, Changsha’s farm has about 8.21 million live birds in stock, with an average daily production of more than 90000 live birds, of which more than 20000 are sold abroad, more than 50000 are sold in the live poultry trading market, and more than 20000 are sold directly to the retail market. The wholesale market stock is 4525, with daily purchases and sales of more than 130000, of which about 80000 live birds come from other places. The retail market has a stock of 30318, with daily purchases and sales of more than 150000, of which 130000 come from the wholesale market and 20000 from farms. The city has a population of 7.04 million, with an average of 100 people consuming 2 live birds per day.

About 64% of the live poultry were from Changsha and other cities in Hunan Province, and the other 36% were from Hubei Province, Henan Province, Guangxi Province, Jiangxi Province, and Guangdong Province (Figure 1-A). Although 79.01% of the live poultry produced in Hunan Province, about 20.99% of them were traded out to eight provinces (Figure 1-B).

**Figure 1.**
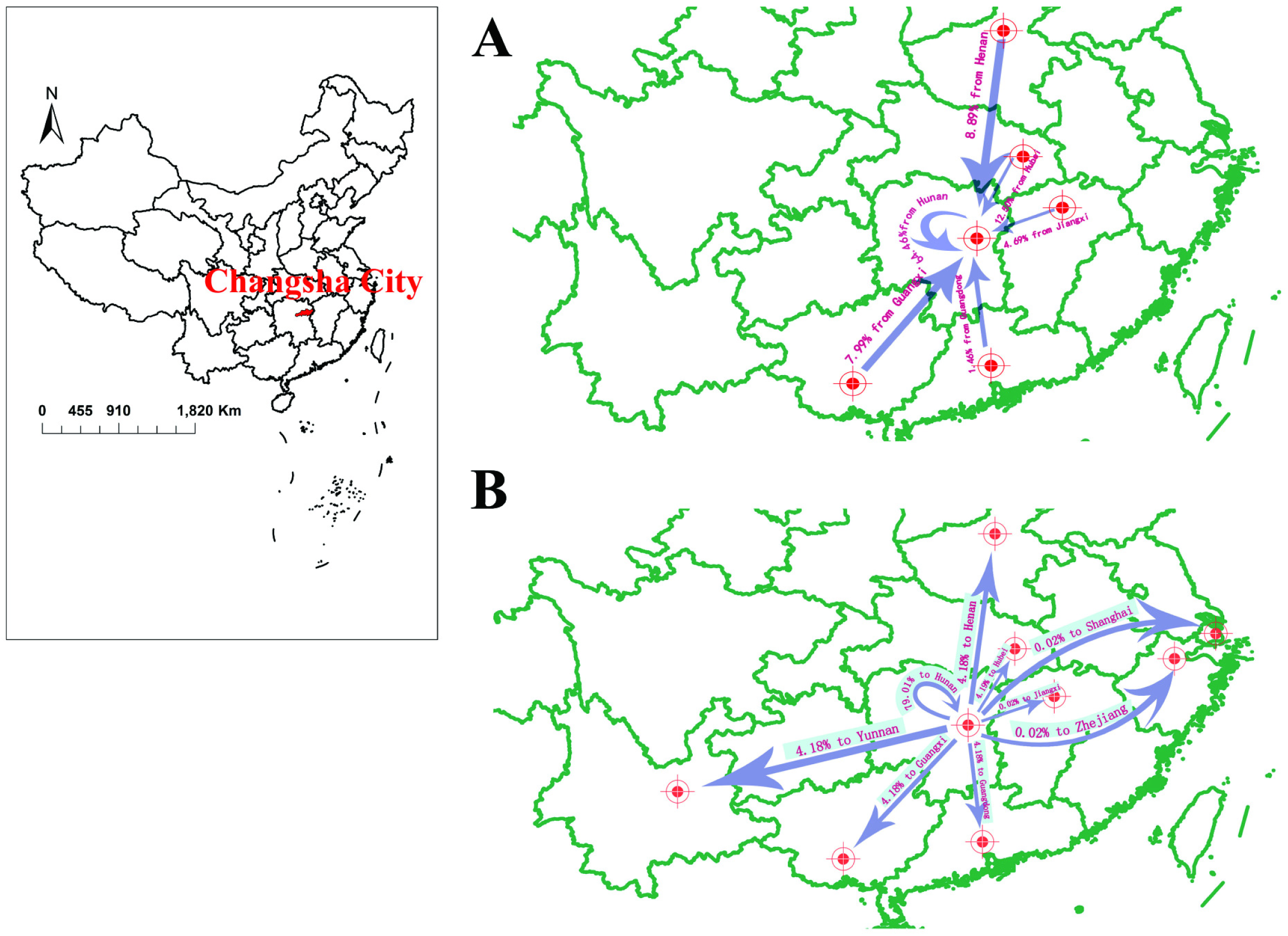
Modes of live poultry from and to places outside Changsha City, China, 2014. A, the sources of live poultry sold in Changsha; B, the live poultry sold to places outside the city.

### Avian virus at the LPFTN network

From January 2014 through March 2015, 840 samples of drinking water, sewage, fecal dropping swabs, and poultry cage swabs were collected from the LPFTN network. RT-PCR results demonstrated that 71 of the 840 analyzed environmental samples (8.45%) were positive for N6. October, November, and July, 2014 had the first highest positive rate. Different sample types had different prevalence of H5N6. The first highest positive rates were observed from chopping board swabs, sewage samples, and drinking water samples (Table 2). From the 71 positive specimens, 10 A(H5N6) viral genomic sequences were obtained for the further analysis.

**Table 2.**
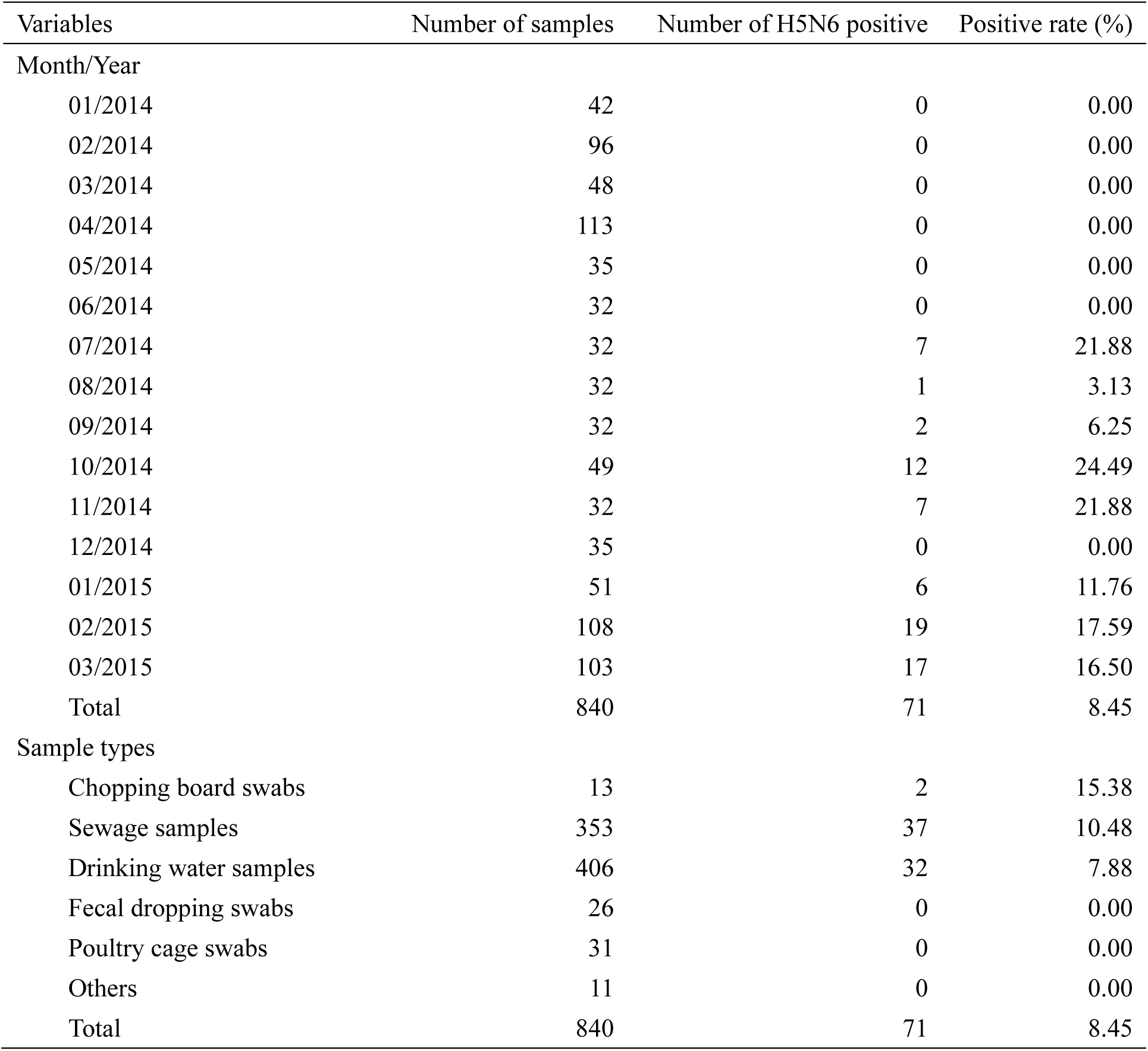
Investigation of H5N6 virus in the LPFTN network in Changsha City, China, 2014

**Table 3.** Nucleotide sequence similarities between the 10 avian influenza A(H5N6) environmental viruses isolated in Changsha, Hunan province, China in 2014 and A/Changsha/1/2014(H5N6) isolated from a patient on February 16, 2014

| CS1* | Collection date | PB2 | PB1 | PA | HA | NP | NA | MP | NS |
| --- | --- | --- | --- | --- | --- | --- | --- | --- | --- |
| CS 308 | 2014 Jun 17 | 98.0 | 98.1 | 98.7 | 92.8 | 97.6 | 87.9 | 98.4 | 98.0 |
| CS 399 | 2014 Sep 18 | 97.6 | 97.8 | 98.6 | 93.0 | 98.2 | 87.6 | 98.4 | 97.8 |
| CS 441 | 2014 Oct 27 | 97.7 | 97.5 | 98.5 | 92.8 | 94.8 | 87.5 | 98.5 | 98.1 |
| CS 443 | 2014 Oct 27 | 97.7 | 97.7 | 98.4 | 92.7 | 98.0 | 87.3 | 98.4 | 98.1 |
| CS 445 | 2014 Oct 27 | 97.6 | 97.7 | 98.3 | 92.6 | 98.0 | 87.6 | 98.3 | 93.0 |
| CS 448 | 2014 Oct 28 | 97.5 | 97.7 | 98.4 | 92.7 | 98.0 | 87.6 | 98.5 | 92.6 |
| CS 455 | 2014 Oct 31 | 97.6 | 98.8 | 98.7 | 92.6 | 98.7 | 87.2 | 98.5 | 98.1 |
| CS 458 | 2014 Oct 31 | 97.4 | 98.7 | 98.7 | 92.7 | 98.7 | 87.4 | 98.3 | 97.7 |
| CS 488 | 2014 Nov 25 | 97.7 | 97.5 | 98.7 | 92.7 | 98.2 | 87.5 | 98.4 | 98.0 |
| CS 499 | 2014 Nov 26 | 97.4 | 97.8 | 98.5 | 92.5 | 97.6 | 87.4 | 98.5 | 94.7 |

\* CS1, A/Changsha/1/2014(H5N6); CS308, A/environment/Changsha/308/2014; CS399, A/environment/Changsha/399/2014; CS441, A/environment/Changsha/441/2014; CS443, A/environment/Changsha/443/2014; CS445, A/environment/Changsha/445/2014; CS448, A/environment/Changsha/448/2014; CS455, A/environment/Changsha/455/2014; CS458, A/environment/Changsha/458/2014; CS488, A/environment/Changsha/488/2014; CS499, A /environment/Changsha/499/2014.

### Avian virus detected by the ISS and PSS systems

From February 17, 2014 to December 1, 2016, a total of 17 laboratory-confirmed cases of human infection with avian influenza A(H5N6) virus in China have been reported by WHO (Table 1)^1^.

In 2014, the ISS system monitored 468041 patients among which 25670 were ILI cases in Changsha City. A total of 2147 throat swabs were collected from the ILI cases to detect the influenza viruses. About 16.63% (357/2147) of the tested cases were confirmed as influenza virus infection. Moreover, there were 189 cases confirmed as H3N2, 101 as influenza B, 66 as influenza A (H1N1), and 1 as H5N6.

The PSS system monitored 798792 patients among which 62726 were pneumonia cases in Changsha City. There were 81 cases suspected as avian influenza infection and their throat swabs or low respiratory tract specimens were collected to detect the influenza viruses. About 45.68% (37/81) of them were tested positive of influenza viruses. Moreover, 21 cases were confirmed as influenza A (H1N1), 5 were H3N2, 4 were influenza B, and no H5N6 were detected by the system in 2014.

### Analysis of the environmental avian influenza A(H5N6) virus strains isolated from LPFTN network

The phylogenetic analyses suggested that most of the 10 environmental A(H5N6) viruses samples obtained in current study from the LPFTN network in Changsha City clustered together (A/environment/Changsha/308/2014, A/environment/Changsha/399/2014, A/environment/Changsha/441/2014, A/environment/Changsha/443/2014, A/environment/Changsha/445/2014, A/environment/Changsha/448/2014, A/environment/Changsha/455/2014, A/environment/Changsha/458/2014, A/environment/Changsha/488/2014, and A/environment/Changsha/499/2014), and have a close relationship to the poultry samples in Vietnam, Nha Trang and Quang Ninh, which demonstrated their avian origin (Figure 2).

**Figure 2.**
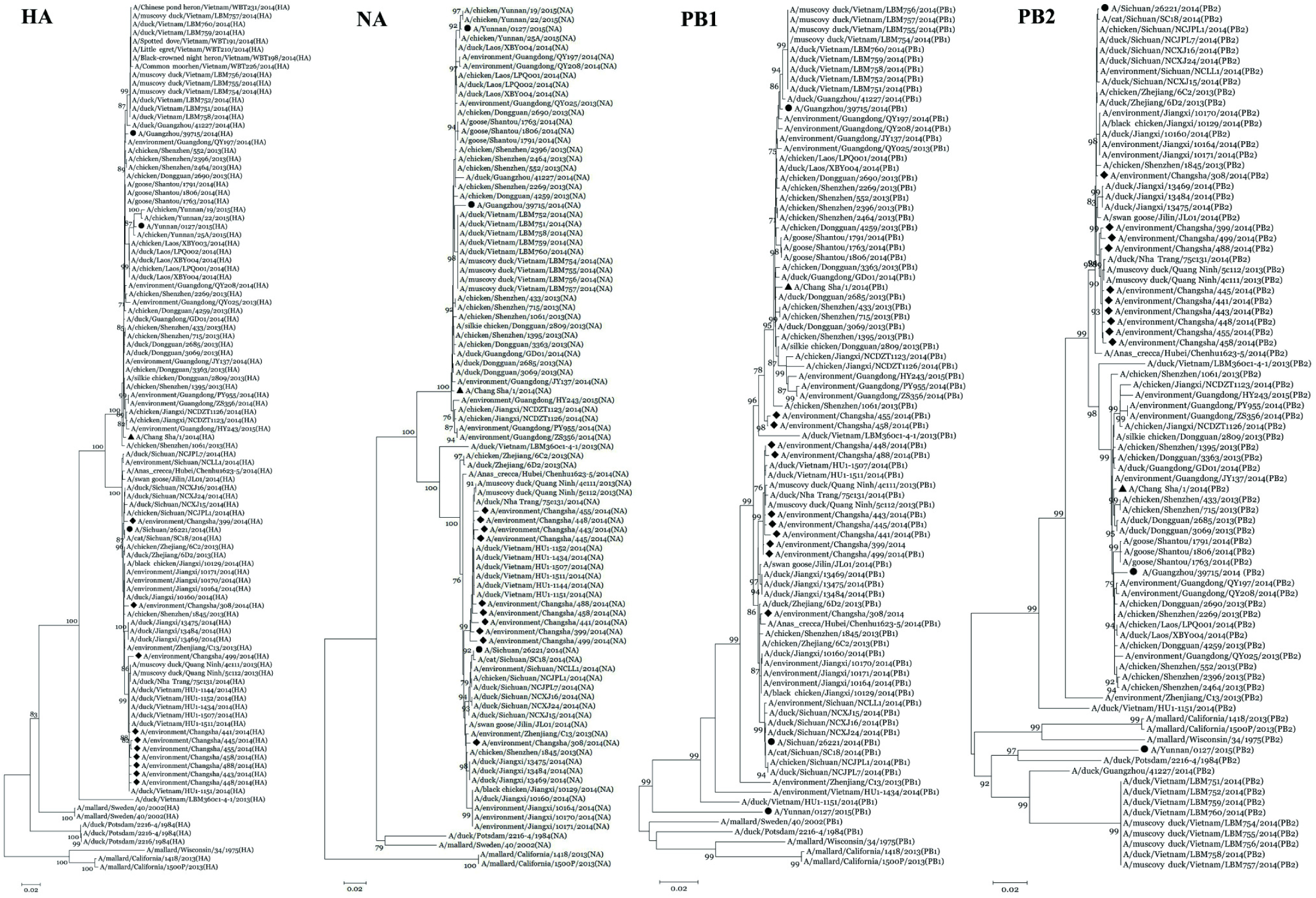
Phylogenetic analysis of the hemagglutinin (HA), neuraminidase (NA), polymerase basic protein (PB1), and PB2 of the novel avian influenza A(H5N6) virus isolated from patients, environment, and poultry. ♦, A(H5N6) viral sequence isolated from an environmental sample; ▴, viral sequence isolated from a patient in Changsha; •, viral sequences isolated from three patients reported in China.

In addition, these environmental samples were shown to be highly similar to the sequences of the local human isolate A/Changsha/1/2014(H5N6), a 5-year-old girl who developed fever and mild respiratory symptoms on February 16, 2014 in Changsha, Hunan province, China^6^: PB2, 97.4%-98.0%; PB1, 97.5%-98.8%; PA, 98.3%-98.7%; HA, 92.5%-93.0%; NP, 97.6%-98.7%; M, 98.3%-98.5%; and NS, 92.6%-98.1%, with a moderate level of similarity between the NA gene sequences (87.2%-87.9%) (Table 3).

### Phylogenetic analyses of avian influenza A(H5N6) virus isolated from patients

Phylogenetic analyses revealed that the patient strain A/Chang Sha/1/2014(H5N6) was closely related to the poultry and environmental A(H5N6) isolates obtained from local and the LPMs in Guangdong province, as well as the poultry samples collected in Jiangxi province, China (Figure 2). Specifically, nucleotide identity of the RNA polymerase basic protein 2 (PB2) gene segments obtained from the A/Chang Sha/1/2014(H5N6) isolate was shown to be similar to that of A/Guangzhou/39715/2014(H5N6) virus and A/environment/Guangdong/JY137/2014(H5N6) isolate (both 99.8%). RNA polymerase basic subunit 1 (PB1) and Hemagglutinin (HA) gene segments were closely related to A/duck/Dongguan/3069/2013(H5N6) (99.6%) and A/chicken/Dongguan/3363/2013(H5N6) (99.5%). Both the gene segments neuraminidase (NA) and Matrix protein (MP) showed the highest nucleotide identity of 99.7% to A/duck/Dongguan/2685/2013(H5N6). While the polymerase acidic protein (PA) was closely related to that of A/environment/Guangdong/JY137/2014(H5N6) (99.7%). And the nucleocapsid protein (NP) and nonstructural protein (NS) were shown to be closely related to that of A/duck/Guangzhou/018/2014(H5N6) with the similarity of 99.7% and 99.8, respectively (Figure 3).

**Figure 3.**
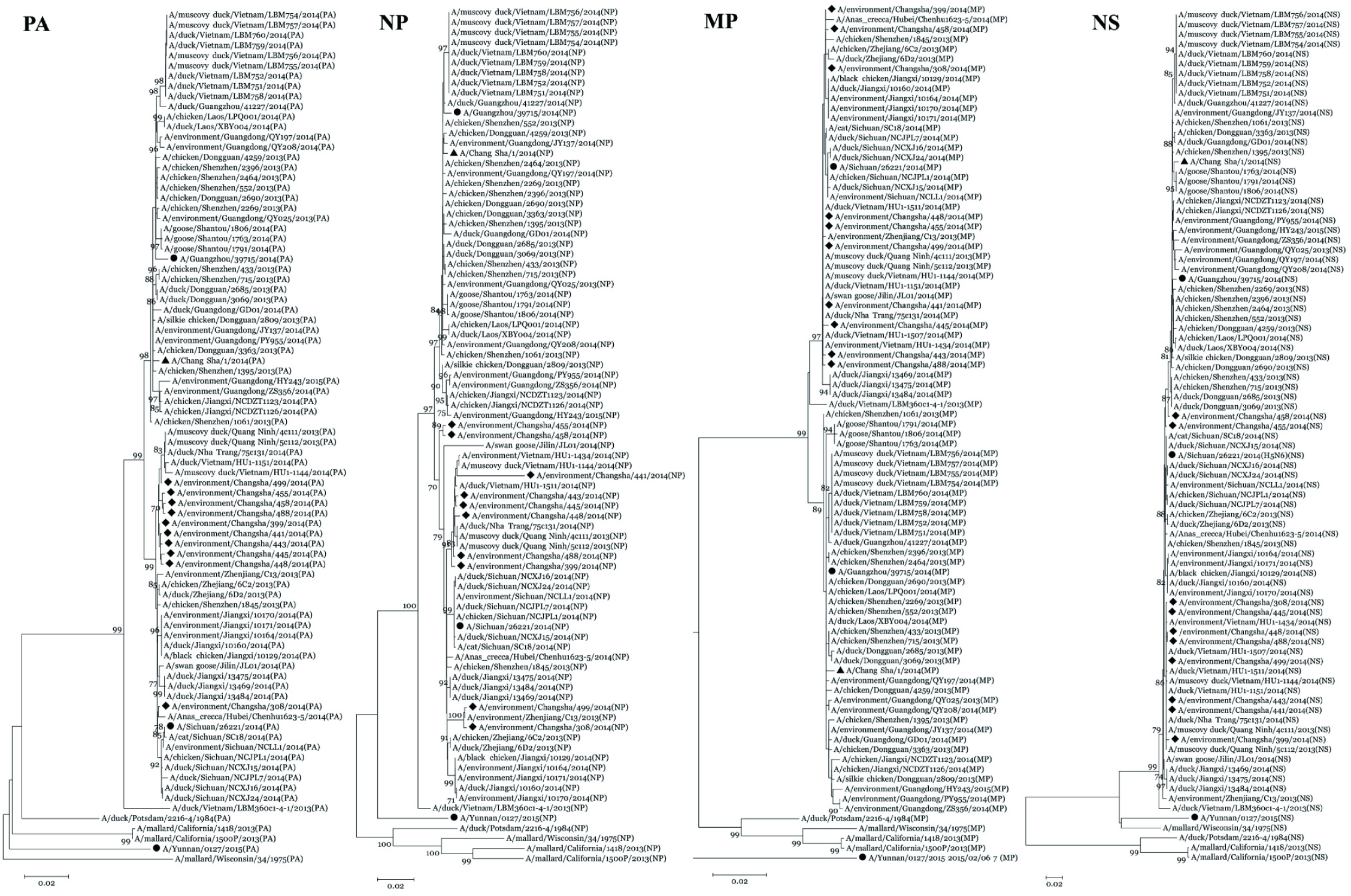
Phylogenetic analysis of the polymerase acidic (PA), nucleocapsid protein (NP), matrix protein (MP), and nonstructural protein (NS) of the novel avian influenza A(H5N6) virus isolated from patients, environment, and poultry. ♦, A(H5N6) viral sequence isolated from an environmental sample; ▴, viral sequence isolated from a patient in Changsha; •, viral sequences isolated from three patients reported in China.

Furthermore, we analyzed A/Sichuan/26221/2014(H5N6) virus strain, isolated from a 49-year-old male who developed clinical symptoms on April 13, 2014 in Nanchong, Sichuan province, China^3^, and phylogenetic analyses revealed that it was closely related to the isolates detected in cat, poultry, and environmental samples from the LPMs in Sichuan province (Figures 2 – 3). Nucleotide identities of the PB2, PB1, PA, and NA genes were shown to be closely related to those detected in A/cat/Sichuan/SC18/2014(H5N6) (99.8–99.9%), A/chicken/Sichuan/NCJPL1/2014(H5N6) (99.7–99.9%), and A/environment/Sichuan/NCLL1/2014(H5N6) (99.9%) isolates, respectively. Nucleotide identities of the HA and NP genes were shown to be highly similar to those identified in A/cat/Sichuan/SC18/2014(H5N6) (99.9–100.0%), A/chicken/Sichuan/NCJPL1/2014(H5N6) (99.8–100.0%), and A/duck/Sichuan/NCXJ16/2014(H5N6) (99.8–99.9%) isolates, respectively. Furthermore, nucleotide identities of the MP gene and NS genes in the virus strain were shown to be the identical to A/chicken/Sichuan/NCJPL1/2014(H5N6) and A/duck/Sichuan/NCXJ15/2014(H5N6) isolate, respectively.

Phylogenetic analyses of virus strain A/Guangzhou/39715/2014(H5N6), isolated from a 59-year-old male registered on December 4, 2014 in Guangzhou, Guangdong province, China^5^, showed that it was closely related to the poultry and LPM environmental isolates from Guangdong and poultry isolates from Laos (Figures 2 – 3). For this strain, the nucleotide identity of the PB2 was shown to be close to those of A/environment/Guangdong/QY197/2014(H5N6) (99.0%) and A/chicken/Shenzhen/2269/2013(H5N6) (99.1%), while PB1 and HA matched those identified in the virus strain A/environment/Guangdong/QY197/2014(H5N6) (99.6% similarity), and the remaining genes all best matched to the poultry samples with high similarities: the PA gene matched the sequence obtained from A/chicken/Shenzhen/2464/2013(H5N6) virus strain (99.3%), NP sequence matched the same gene in the virus strain A/duck/Guangzhou/41227/2014(H5N6) (99.5%), NA sequence was closely related to that detected in A/chicken/Shenzhen/2269/2013(H5N6) (98.8%), while MP and NS sequences were shown to be similar to those of A/chicken/Laos/LPQ001/2014(H5N6) (100%) and A/chicken/Shenzhen/552/2013(H5N6) (99.5%), respectively.

We further analyzed A/Yunnan/0127/2015(H5N6) isolate, obtained from a 44-year-old male who developed avian influenza A symptoms on February 8, 2015 in Shangri-La City, Yunnan province, China ^4^, and demonstrated that the HA and NA genes in this strain closely resembled those of the A(H5N6) viral strains isolated from chicken in Yunnan province (Figures 2 – 3), and the HA and NA gene sequences were similar to those of the A/chicken/Yunnan/25A/2015(H5N6) (99.6%) and A/environment/Yunnan/YN19/2015(H5N6) (99.9%) genes, respectively. The sequences of 6 genes of the A/Yunnan/0127/2015(H5N6) virus were shown to have a high level of similarity to those of avian influenza A(H9N2) and (H7N9) viruses (98.2-99.6%), isolated from birds and humans in China.

### Analysis of avian influenza A(H5N6) viral protein sites in strains isolated from patients and the LPFTN network

Although A(H5N6) viruses may be highly pathogenic in poultry ^16^, we did not detect 6 basic amino acid (R, K) mutations at the cleavage site and the substitution of Q226L and G228S at the 210-loop in the HA protein. These mutations, when present in the viral strains isolated from humans and environment (Tables 4 – 5), may facilitate human infections ^17^.

**Table 4.**
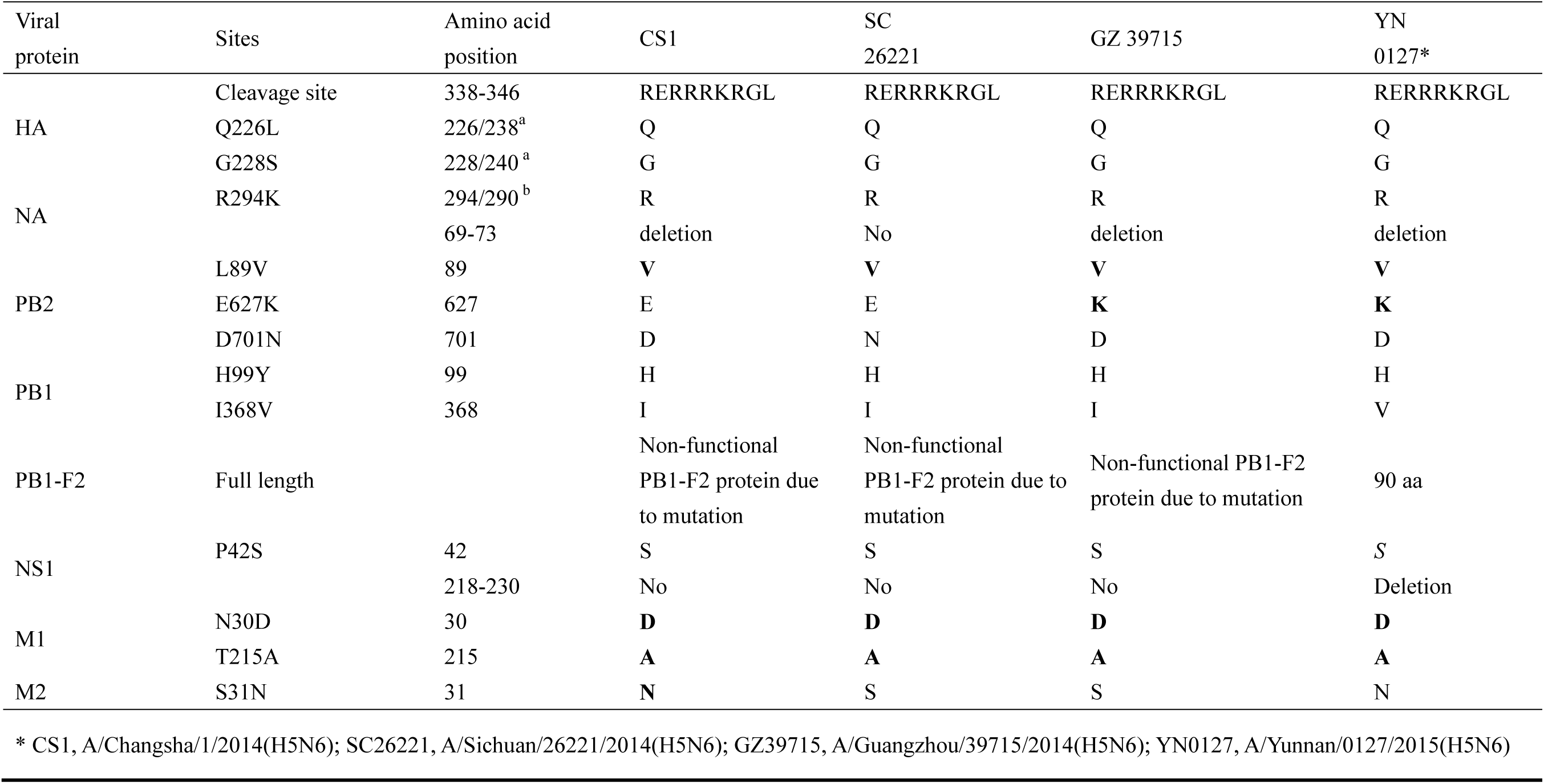
Analysis of the eight viral proteins of the four human avian influenza A(H5N6) viral strains

**Table 5.**
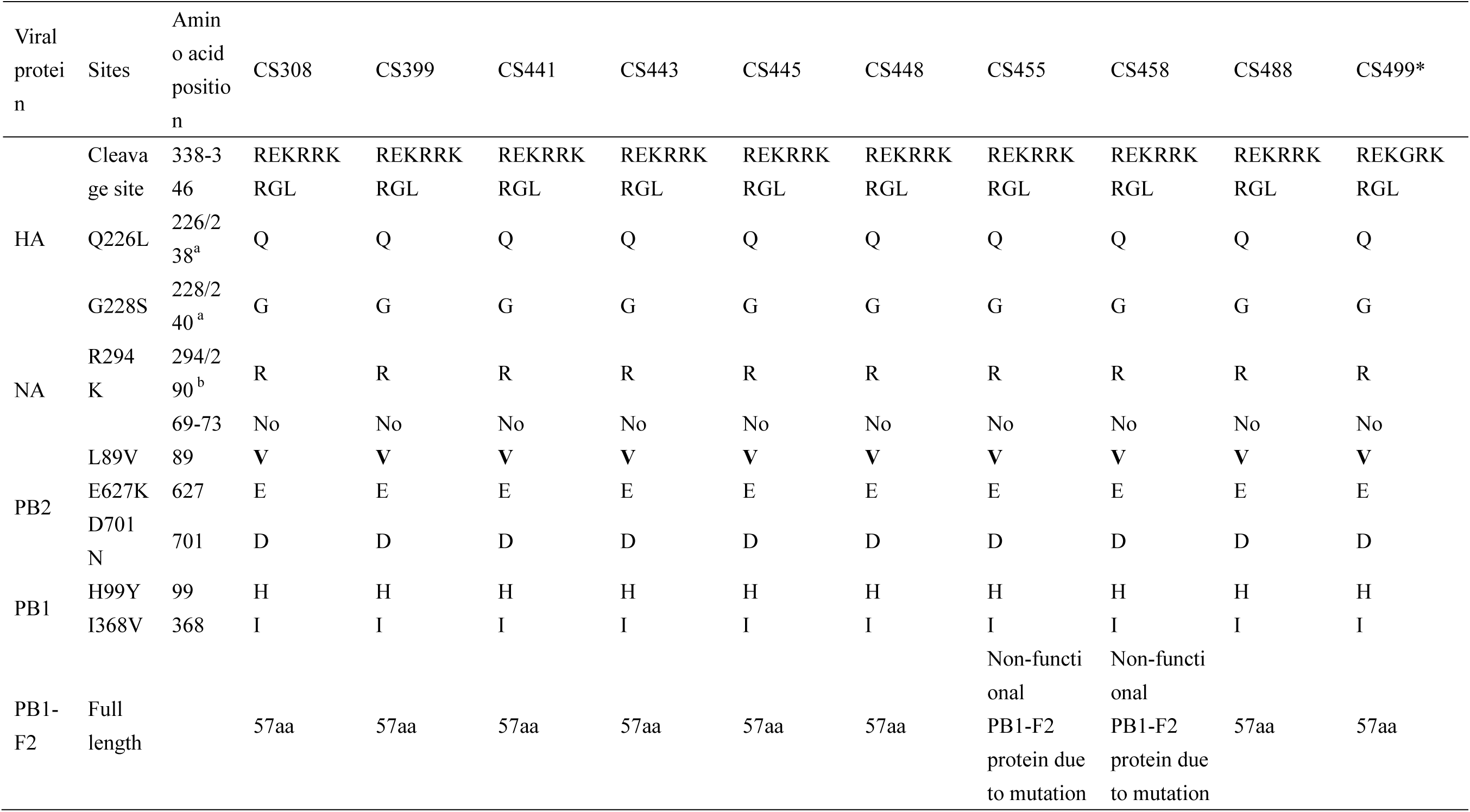
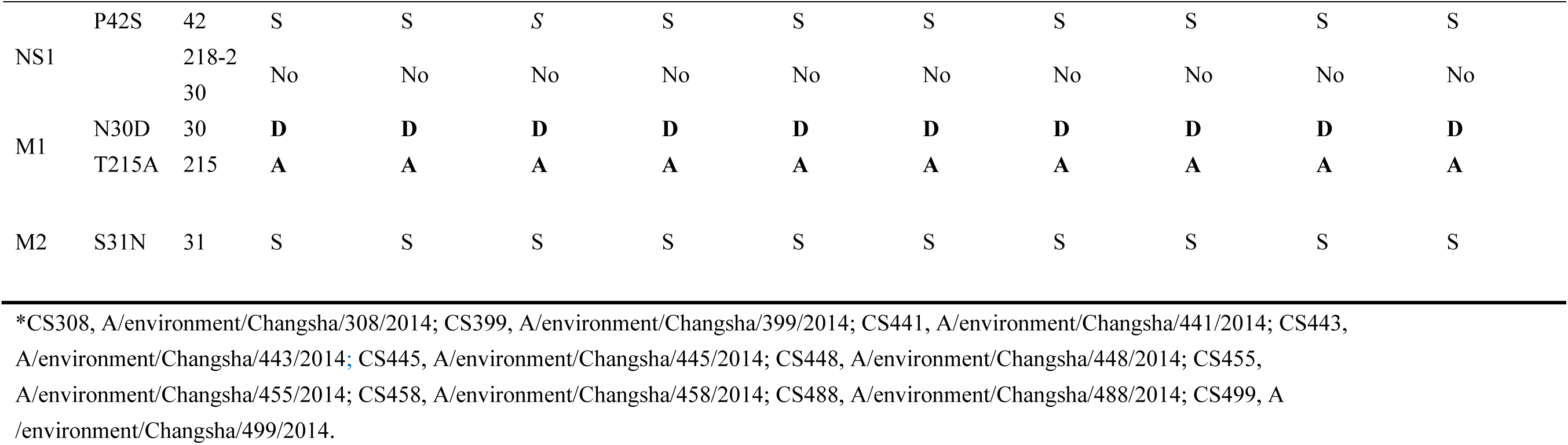
Analysis of the eight viral proteins of the 10 environmental avian influenza A(H5N6) viral strains

Resistance-conferring mutation, R294K in NA, was not observed in the human and environmental isolates, indicating that these viruses are sensitive to neuraminidase inhibitors ^18^. However, the substitution S31N in the M2 gene was observed in two viral strains isolated from patients, A/Changsha/1/2014 and A/Yunnan/0127/2015, which confers the resistance to adamantanes (Table 4)^19, 20^, but it was not found in other analyzed viruses. In 3 A(H5N6) viral stains isolated from patients, A/Changsha/1/2014, A/Guangzhou/39715/2014, and A/Yunnan/0127/2015, a deletion in the stalk region of NA, residues 59 to 69, was detected, but this was not observed in other human and environmental isolates (Tables 4 – 5). E627K, another mutation in PB2 that facilitates the adaption to mammalian hosts ^21^, was observed in 2 viral strains obtained from patients (A/Guangzhou/39715/2014 and A/Yunnan/0127/2015).

## Discussion

In this study, we demonstrated that the LPMs represent a major source of human infection with the avian influenza A(H5N6) virus in China, and in many of these, different subtypes of this virus can be found. In a previous study, a correlation between the A(H5N6) virus isolated from a patient in Changsha and those isolated from the local LPMs that patient had visited prior to developing the infection, was not found ^6^. In this study, we expanded the number of samples collected from several LPMs and demonstrated that large amounts of A(H5N6) viruses can be detected in the LPM environment in Changsha. The sequence of the NA gene of the viral strain isolated from a patient, A/Changsha/1/2014, from Changsha, was shown to be closely related to those of the A(H5N6) viral strains isolated from the LPMs in Changsha. Additionally, these viruses were isolated from the LPM samples on February 15, 2014, which is similar to the time of the Changsha patient’s exposure to this virus.

Epidemiological investigation in a case reported in Sichuan demonstrated that the source of A(H5N6) virus was a local LPM, but this virus was not detected at that market^3^. In our study, we collected the data on all reported A(H5N6) viral infections from the NCBI database, and our phylogenetic analyses revealed that the human A(H5N6) viral isolate (A/Sichuan/26221/2014) was closely related to viral strains isolated from cat, poultry, and environmental samples collected from the LPMs in Sichuan province. These results indicate that the infected poultry sold at the local LPM was the source of human infections in Sichuan. Furthermore, by analyzing the origin of infection of a patient in Guangdong, it was demonstrated that the patient had visited a LPM before the onset of disease ^5^. Phylogenetic analyses revealed that the viral strain isolated from this individual was closely related to those isolated from poultry and LPMs in the same province, which further suggests that the LPMs are a major source of human infections with avian influenza A in Chinese provinces.

Currently, 2 human infection cases with A(H5N6) virus have been reported in Shangri-La City, Yunnan province, China, but the sequence of only one of these strains has been published in the NCBI database. One of these patients was exposed to the dead wild fowl, while the exposure history of the other, confirmed to be infected with avian influenza A(H5N6) virus accompanied by severe pneumonia ^22^, is unknown. Our investigations revealed that the HA and NA gene sequences detected in the virus isolated from one of the described patients, A/Yunnan/0127/2015, were closely related to that of the A/chicken/Yunnan/25A/2015(H5N6) virus strain, isolated from the poultry in Yunnan province. Therefore, our phylogenetic analyses and sequence comparisons suggest that the poultry and/or LPMs represent the major sources of human infection with avian influenza A(H5N6) virus in China.

The recent outbreaks of the A(H5N6) viral infections in poultry, wild birds, and domesticated animals in southeastern Asia ^7-9, 23, 24^ demonstrate the diversity of the sources of A(H5N6) virus gene reassortments ^2, 6, 10^. In China, the purchasing of live poultry in the LPMs is a common practice, and the infections of humans with other subtypes of viruses at the LPMs have been previously confirmed ^11-13^. The results of our study demonstrate that the surveillance of the avian influenza A viruses at the LPMs throughout China should be increased. Furthermore, we believe that the results of our study may help understand the importance of the LPM cleaning, and changing and regulating the modes of live poultry sale.

